# The critical role of HDAC1 activates NSCLC growth by nicotine resistance Cisplatin

**DOI:** 10.1101/2020.06.04.992347

**Authors:** Ching-Yi Peng, Jia-Ping Wu

## Abstract

Nicotine is active in highly cisplatin-resistant cancer cells; however, there is little evidence for its resistant activity in lung cancer with cisplatin. Many mechanisms of cisplatin resistance have been proposed. The mechanisms of the nicotine treatment of cisplatin-resistant lung cancer for histone deacetylase 1 (HDAC1) activity is unknown. Nicotine was used to analyze cisplatin-resistant non-small cell lung cancer (NSCLC) cancer cell growth. Western blot was used to analyze cell cycle-related proteins. Cancer cell viability (cell survival) was measured with MTT assay. HDAC1 transfected NSCLC cells were used to analyze the direct binding between cytosol and nucleus distribution. Here, using cell viability and migration methods we firstly found nicotine regulated cisplatin-resistant NSCLC cells growth by targeting HDAC1. Expression of cisplatin was negatively correlated with HDAC1. And HDAC1 inhibitor, VPA, in the NSCLC cancer cells were predicted. Further experiments confirmed that HDAC1 directly targeted E2F and cisplatin. Besides, HDAC1 and cisplatin inhibited NSCLC cell growth and reduced expression of E2F and Cyclin E proteins. The use of nicotine compromised cisplatin-induced E2F suppression and cancer cell growth. NSCLC cancer cells co-transfected with nicotine and HDAC1 had a higher cell cycle proliferation. Taken all together, cisplatin interferes with DNA replication kills the cancer cell fastest proliferation; however, nicotine increased detoxification of cisplatin, inhibition of apoptosis and DNA repair, induced cisplatin resistance.

## Introduction

Cigarette smoke exposure promoted NSCLC lung cancer in all patients, because of the role of cigarette smoke extract, nicotine, exhibits promotion co-carcinogenesis and activities in tumor production and malignant transformation [1,2]. Cigarette smoke extract, nicotine, induces promotion co-carcinogenesis may have crucial roles with cancer growth remains obscure. In contract, the underlying mechanisms nicotine through α7nAChR nicotine receptor or EGFR to resistant traditional chemotherapies like cisplatin in lung cancer is still unknown. Nicotine constitutes of the dry weight of cigarette [3]. Once exposure enters the body, nicotine not only associated with cancer in humans, but also metabolized [4]. It is highly addictive to NSCLC lung cancer. Lung Cancer development may also involve epigenetic changes [5,6]. The main epigenetic modifications in humans are histone modifications. Histone deacetylases enzymes (HDAC) is an enzyme to lead to histone chemical modification after translation [7]. The acetylation of histones level controlled post-translation modification and epigenetic gene regulation. HDAC plays a circle role in cancer physiological roles including development growth, depending on the specific extracellular environment [8]. Histone deacetylase inhibitor, valproic acid (VPA), is a short-chain fatty acid that can cause growth arrest and induce differentiation of transformed cells [9-11]. Cisplatin chemotherapy is the cornerstone of treatment of lung cancer. Cisplatin is a well-known chemotherapeutic drug. Cisplatin interferes with DNA replication kills the fastest proliferating cancerous. The main function of cisplatin is to inhibit the different phases of the process of forming the tumor cell cycle to kill cancer cells to treat tumors [12,13]. Initial platinum responsiveness is high but the majority of cancer patients will eventually relapse with cisplatin-resistant disease. Cigarette smoke exposure stimulates angiogenesis and tumor growth increased resulted in detoxification of cisplatin, inhibition of apoptosis and increased cancer cell DNA repair. Cigarette smoke extract, nicotine, is unusual in comparison to most drugs, as its concentration profile changes from induced tumor growth to cytotoxic with increasing doses [14]. The concentration of nicotine stimulated cell growth to correspond to low concentration was needed, while high concentration was cytotoxic. Many mechanisms of cisplatin-resistance disease have proposed including changes in cellular uptake and efflux of nicotine [15]. Nicotine may be useful in the treatment of cisplatin-resistant cancer. Cisplatin maybe combine to attack cancer cells in bad environment, for example, smoky or nicotine exposure environment always accelerate lung cancer diseases led to chemotherapy drugs lose their functions to kill cancer cells [16]. Nicotine may have an important role to regulate chemotherapy drugs functions. There are many mechanisms interacted with nicotine and chemotherapy drugs.

## Results

### Without nicotine, the interaction of HDAC1 accounts for a chemotherapy drug, cisplatin, in NSCLC cells

The underlying mechanisms of the experimental works described the interaction of HDAC1-related signaling with carcinogenic content cisplatin or HDAC1 inhibitor, valproic acid (VPA) in A549 cells. To develop interaction of HDAC1-related cisplatin treatment strategies for the interplay HDAC1 without nicotine as well as potentially resulted in cisplatin treatment resistance in adenocarcinoma A549 cell. We detected adenocarcinoma A549 cell viability (%) after treatment transfected HDAC1^+/+^ plasmid, valproic acid (VPA), cisplatin (Cis) and combining VPA and Cis at 24 and 48 h using MTT analysis. Results showed transfected HDAC1^+/+^ plasmid has significantly increased cell viability (%) at 48 h (p<0.05). Valproic acid (VPA), cisplatin (Cis) and combining VPA and Cis treated adenocarcinoma A549 cell viability were lower than control at 24 and 48 h treatment (p<0.01). VPA led to adenocarcinoma A549 cell viability at 48 h lower than 24 h (p<0.01). It should be noted that cisplatin (Cis) at 24 h treated A549 cell has significantly decreased cell viability (%) compared with at 48 h (p<0.05) (Fig. 1A and Fig. S1A). Chemotherapy drugs, cisplatin, dose at 25 and 10 μM has significant decreased by MTT analysis cell viability (%) (p<0.05) (Fig. 1B and Fig. S1A). Therefore, we used this dose as following examination. Treatment with valporic acid (VPA), a HDAC2-specific inhibitor, enhanced the adenocarcinoma A549 cell viability cytotoxicity at 24 h using MTT analysis. Valporic acid (VPA) dose at 50 and 100 mM has significantly decreased cell viability (%) at 24 h (Fig. 1C and Fig. S1A). Contractually, different levels of HDAC1 protein expression in transfected HDAC1 plasmid (HDAC1^+/+^), siHDAC1 and VPA treatment without nicotine at 24 h using immunofluorescent staining. Result shown HDAC1^+/+^ plasmid transfected has higher levels than control, in contrast, siHDAC1 and VPA treatment without nicotine has lower in adenocarcinoma A549 cells by immunofluorescence staining (Fig. 1D and Fig. S1B). From cancer cell move sooner or not, we found HDAC1^+/+^ plasmid transfected has the greatest expression, but the others were not observed by migration (Fig. 1E). To compare the impacts of the HDAC1-related on HDAC1^+/+^ plasmid transfected, cisplatin, and VPA without nicotine in NSCLC cancer cells will be freshly produced by the following indicated cell cycle arrest, apoptosis induction and activation of cancer-suppressor genes using western blotting analysis following Fig. 2, 3 and 4. Our preliminary immunofluorescent staining results showed HDAC1 expression levels in A549 adenocarcinoma cells after HDAC1^+/+^ transfected plasmid was observed increases, but cisplatin and VPA without nicotine in adenocarcinoma A549 cells were decreases using immunofluorescence staining (Fig. 2A). VPA is an inhibitor of HDACs. Inhibition of HDACs led to cell cycle arrest at G1/S phase by HDAC1 inhibitor. Therefore, we found not only G1/S phase, Cyclin D1/Cyclin E/Cyclin A/E2F protein expression levels increased by HDAC1 plasmid (HDAC1^+/+^), but also increased by HDAC1/HDAC2/HDAC3/HDAC4 (Fig. 2B and 2D). In contrast, VPA, Cis, and VPA combined Cis (VPA+Cis) inhibited Cyclin D1/Cyclin E/Cyclin A/E2F protein expression compared with control (Fig. 2B). Down-regulation of NF-κB and Caspase-3 expression mediates HDAC1 plasmid (HDAC1^+/+^) transfection into A549 cell. NF-κB, Caspase 3, p21, p53, Bax and γ-H2AX protein expression levels was increased by VPA, Cis and VPA+Cis (Fig. 2C and 2D). To detect HDAC activity maybe for developing HDAC-targeting cisplatin drug. HDAC1 plays a circle role in lung cancer cells physiological roles. Therefore, we established HDAC1 knockdown cell lines, siRNA lentivirus plasmid-A, transfection into lung cancer cells for 24 h. From Fig. 2E results, we found cell cycle related protein (cyclin E), Cyclin-dependent kinase inhibitor (p27) and radiation related protein (γ-H2AX) have associated with HDAC1 regulation. Cisplatin also has the same function with HDAC1 (Fig. 2E). Cell cycle suppressors, cyclin-dependent kinase inhibitors (p21, p27 and p53) are associated with linking cell cycle arrest (Fig. 3). Therefore, we found protein expression levels of p21, p27 and p53 by western blotting were decreased by HDAC1^+/+^ transfection, but increased by VPA, Cis and combining VPA and Cis (p<0.05) (Fig. 3A, 3B, and 3C). Cell growth inhibition and apoptosis may be a result of HDAC1-mediated cell cycle suppression. ATM protein was observed by VPA (Fig. 3D). VPA, Cis and combining VPA and Cis were induced BAX expression increased (p<0.05) (Fig. 3E). HDAC1 and NF-κB increases were observed by HDAC1^+/+^ transfection by immunofluorescence staining analysis (Fig. 3F). In addition, we found that HDAC1 location at cell nucleus, but not cytosol. We can from Fig. 3G results to determine it in A549 cell. Long-term 48 h, we observed cisplatin-resistance effects on HDAC1, E2F, Rb and pRb (Fig. 4A). This effect was not observed at 24 h in A549 cell. This is a natural effect cisplatin-resistance at 48 h. On sequamous carcinoma cell, H520, we can find E2F, Rb and pRb have cisplatin-resistance effect at 48 h, not significantly increased. In contrast, p53, p21, p27 and caspase 3 also have cisplatin-resistance, not significantly decreased (Fig. 4B and 4D). HDAC1 also has cisplatin function suppressed H520 cell growth (Fig. 4C). Thus, we determine nucleus and cytoplasm HDAC1 effects and their interaction with cell cycle. According to Fig. 5A result, we found HDAC1 in the nucleus, but not cytosol. HDAC1 plays a circle role in cancer cells physiological roles including development and growth, depending on the specific extracellular environment. We identify the interplay of HDAC1 with cigarette carcinogens (*i*.*e*., nicotine) and potentially resulted treatment cisplatin-resistance in NSCLC in particular adenocarcinoma and squamous cell carcinoma. Nicotine exposure may affect either adenocarcinoma or squamous. After nicotine effects, HDAC1 was from nucleus to cytosol, and released cisplatin binding DNA (Fig. 5A). Notably, nicotine maybe have regulated cisplatin-resistance functions to accelerate lung cancer diseases led to cisplatin lose their functions to kill cancer cells.

**Fig. 1.**
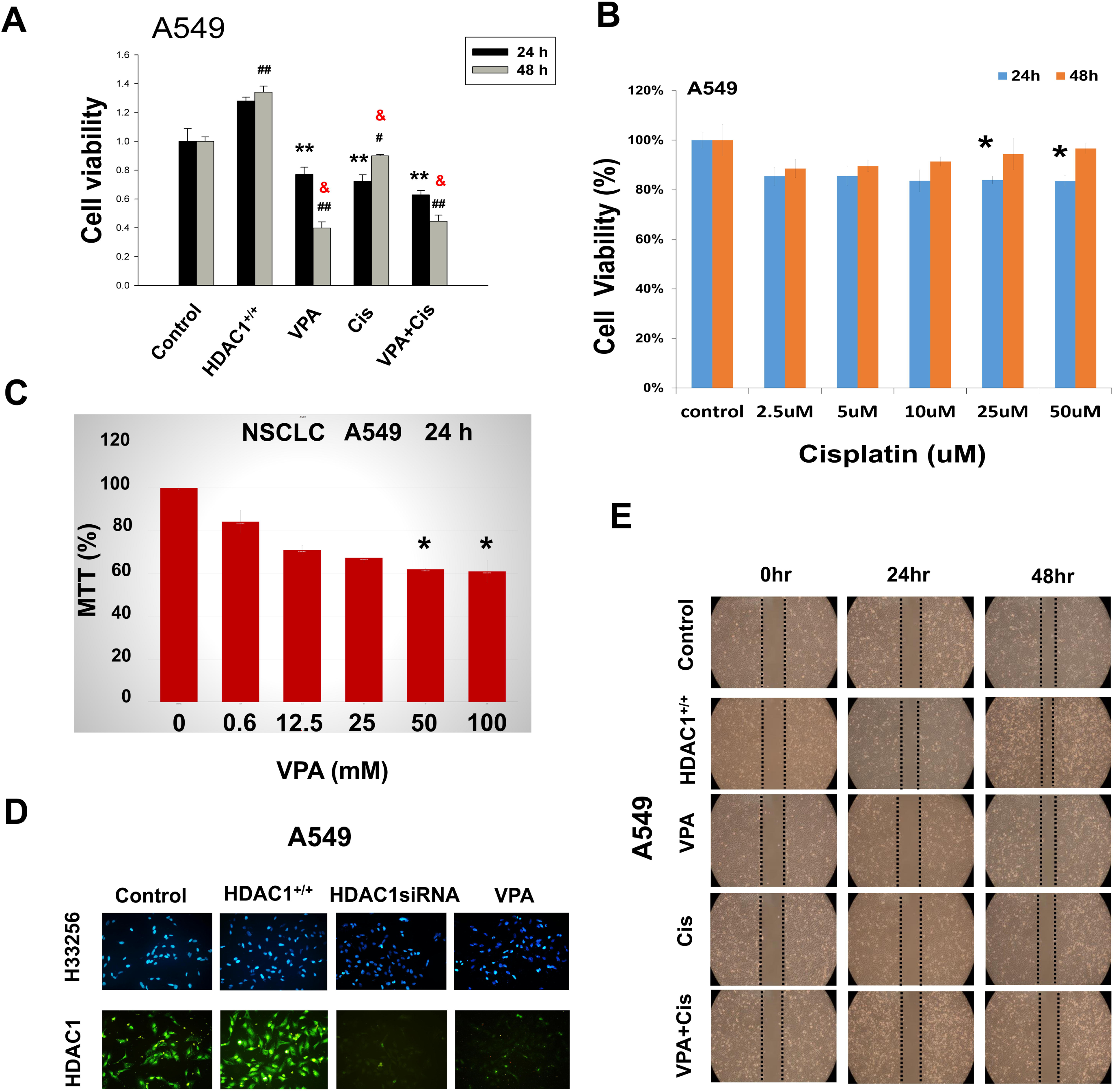
The experimental works described related to carcinogenic content in A549 cells. (A). Cell viability (%) by HDAC1^+/+^, valproic acid (VPA), cisplatin (Cis) and combination VPA and Cis in A549 cell. Statistical analysis of cell viability (%) at 24 and 48 h. Data was expressed mean ± SEM, ^**^p<0.01 significant difference compared with 24 h treatment control. ^#^p<0.05, ^##^p<0.01 significant difference compared with 48 h treatment control. ^&^p<0.01 significant difference compared between with 24 h and 48 h treatment. (B) Cisplatin dose-dependent in A549 cells using MTT assay. Statistical analysis of cell viability (%) in cisplatin treatment A549 cells at 24 and 48 h. Data was expressed mean ± SEM, ^*^p<0.01 significant difference compared with 24 h treatment control. (C) Valproic acid (VPA) dose-dependent in A549 cells using MTT assay. Statistical analysis of cell viability (%) in valproic acid (VPA) treatment A549 cells at 24 and 48 h. Data was expressed mean ± SEM, *p<0.01 significant difference compared with 24 h treatment control. (D) HDAC1 protein expression without nicotine in the cytosol after HDAC1^+/+^, HDACsiRNA, and VPA using immunofluorescent staining. HDAC1^+/+^, HDAC1 plasmid transfection; HDAC1siRNA, HDAC1 small interfering; VPA: valproic acid. Blue color is the nucleus (H33256). Green color is HDAC1 protein expression leves (HDAC1). (E). A549 cells migration at 24 and 48 h after HDAC1^+/+^, VPA, Cis and VPA+Cis. HDAC1^+/+^, HDAC1 plasmid transfection; VPA, valproic acid; Cis, cisplatin.

**Fig. 2.**
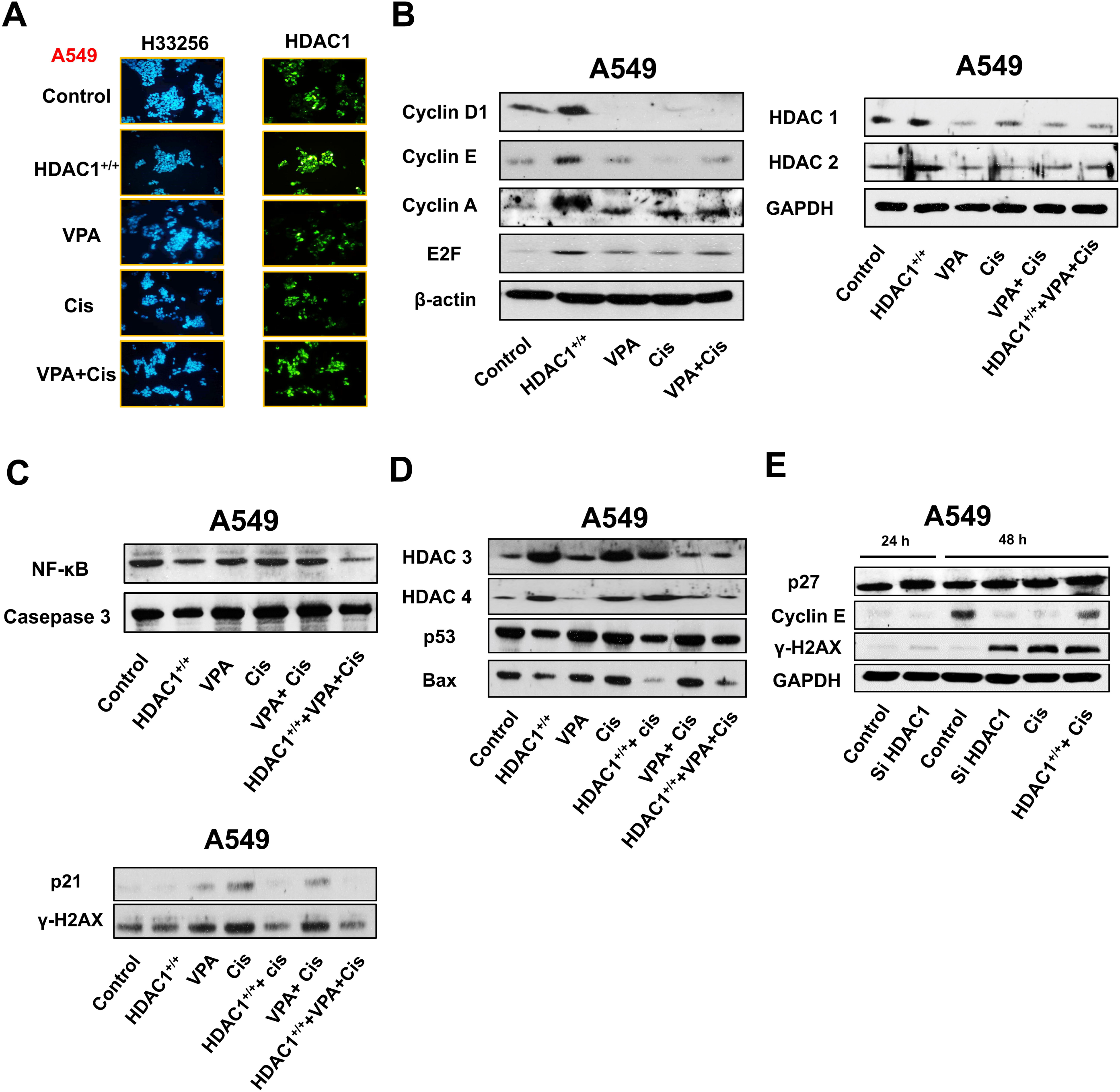
Cisplatin induces an arrest of the cell cycle-related with HDAC1 in A549 cells. (A). By immunofluorescent staining, HDAC1 protein expression without nicotine in the cytosol after HDAC1^+/+^, VPA, Cis and VPA+Cis using immunofluorescent staining. HDAC1^+/+^, HDAC1 plasmid transfection; Cis, cisplatin; VPA, valproic acid. Blue color is the nucleus (H33256). Green color is HDAC1 protein expression leves (HDAC1). (B). Cell cycle checkpoint protein expression of Cyclin D1/Cyclin E/Cyclin A/E2F and histone protein of HDAC1 and HDAC2 using western blotting analysis in A549 cancer cells. (C). Cell apoptosis proteins, NF-kB/p53/p21/γ-H2AX using western blotting analysis in A549 cancer cells. (D). Protein expression of HDAC3/HDAC4/p53/Bax using western blotting analysis in A549 cancer cells. HDAC1^+/+^, HDAC1 plasmid transfection; Cis, cisplatin; VPA, valproic acid. (E). Protein expression of p27/Cyclin E/γ-H2AX using western blotting analysis in A549 cancer cells. HDAC1siRNA, HDAC1 small interfering; Cis, cisplatin.

**Fig. 3.**
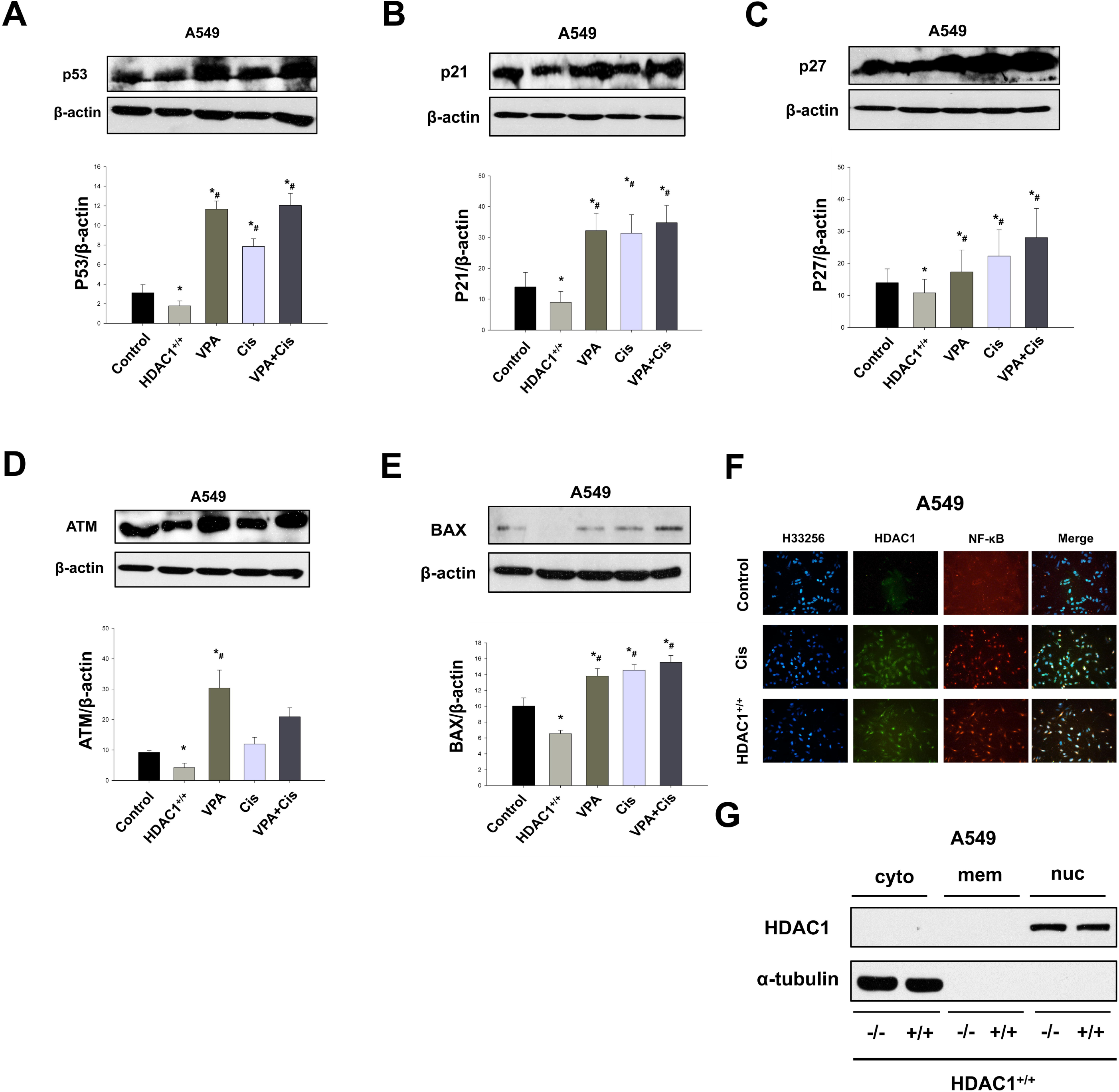
Cell cycle suppression genes related with HDAC1 protein expression in A549 cancer cells after HDAC1^+/+^, VPA, Cis, and VPA+Cis treatment. (A).Cell cycle suppression genes protein expression of p53 using western blotting analysis in A549 cancer cells after HDAC1^+/+^, VPA, Cis and VPA+Cis treatment. Statistical analysis of p53 protein expression. Data was expressed mean ± SEM, * p<0.05, significant difference compared with control. ^#^p<0.05, significant difference compared with HDAC1^+/+^. HDAC1^+/+^, HDAC1 plasmid transfection; Cis, cisplatin; VPA, valproic acid. (B).Cell cycle suppression genes protein expression of p21 using western blotting analysis in A549 cancer cells after HDAC1^+/+^, VPA, Cis and VPA+Cis treatment. Statistical analysis of p21 protein expression. Data was expressed mean ± SEM, * p<0.05, significant difference compared with control. ^#^p<0.05, significant difference compared with HDAC1^+/+^. HDAC1^+/+^, HDAC1 plasmid transfection; Cis, cisplatin; VPA, valproic acid. (C).Cell cycle suppression genes protein expression of p27 using western blotting analysis in A549 cancer cells after HDAC1^+/+^, VPA, Cis and VPA+Cis treatment. Statistical analysis of p27 protein expression. Data was expressed mean ± SEM, *p<0.05, significant difference compared with control. ^#^p<0.05, significant difference compared with HDAC1^+/+^. HDAC1^+/+^, HDAC1 plasmid transfection; Cis, cisplatin; VPA, valproic acid. (D).Cell cycle suppression genes protein expression of ATM using western blotting analysis in A549 cancer cells after HDAC1^+/+^, VPA, Cis and VPA+Cis treatment. Statistical analysis of ATM protein expression. Data was expressed mean ± SEM, *p<0.05, significant difference compared with control. ^#^p<0.05, significant difference compared with HDAC1^+/+^. HDAC1^+/+^, HDAC1 plasmid transfection; Cis, cisplatin; VPA, valproic acid. (E).Cell cycle suppression genes protein expression of BAX using western blotting analysis in A549 cancer cells after HDAC1^+/+^, VPA, Cis and VPA+Cis treatment. Statistical analysis of BAX protein expression. Data was expressed mean ± SEM, * p<0.05, significant difference compared with control. ^#^p<0.05, significant difference compared with HDAC1^+/+^. HDAC1^+/+^, HDAC1 plasmid transfection; Cis, cisplatin; VPA, valproic acid. (F).Immunofluorescent staining of HDAC1 and NF-KB without nicotine in H157 squamous lung cancer cells after HDAC1^+/+^ and Cis treatment. (G).Distributed of HDAC1 in the membrane, cytoplasm and nuclear in A549 cell lines.

**Fig. 4.**
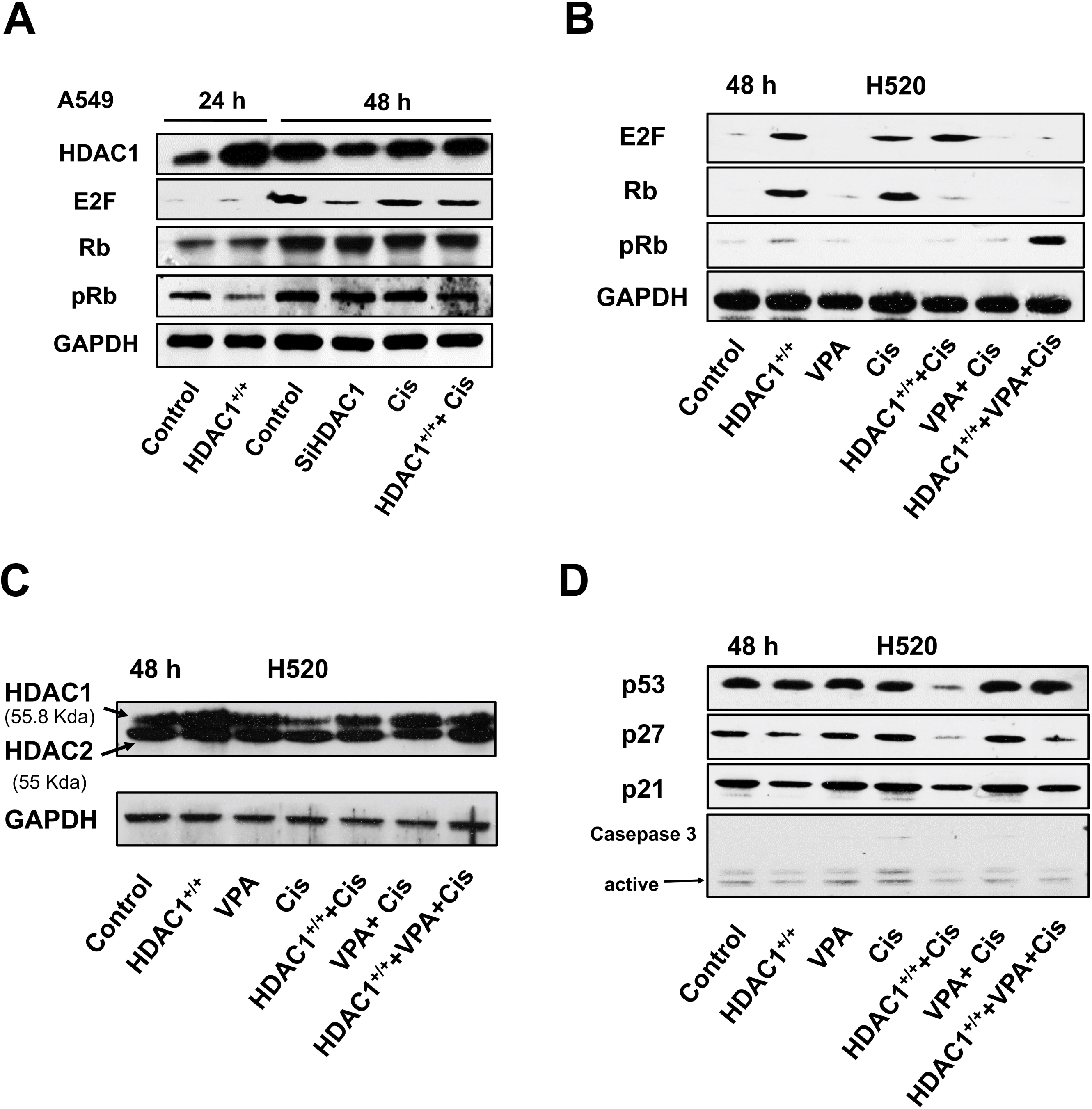
Interaction of HDAC1 and Cisplatin in H520 squamous lung cancer cells. (A).Cell cycle of E2F, Rb and pRb associated with HDAC1 and cis at 24 and 48 h using western blotting analysis in A549 cells. (B).Cell cycle checkpoint proteins, E2F, Rb and pRB associated with HDAC1 and Cis at 48 h using western blotting analysis in H520 cells. HDAC1^+/+^, HDAC1 plasmid transfection; Cis, cisplatin; VPA, valproic acid. (C).The relationship of HDAC1 and HDAC2 with cis at 48 h using western blotting analysis in H520 cells. HDAC1^+/+^, HDAC1 plasmid transfection; Cis, cisplatin; VPA, valproic acid. (D). Cell cycle suppressor genes, p53, p21 and p27 and apoptosis, caspase 3, associated with HDAC1 and Cis at 48 h using western blotting analysis in H520 cells. HDAC1^+/+^, HDAC1 plasmid transfection; Cis, cisplatin; VPA, valproic acid.

### Nicotine exposure induced cisplatin-resistance oncogenic activity not only in adenocarcinoma but also in squamous carcinoma cells

This drug maybe combine to attack cancer cells in different ways. There are many mechanisms interacted with nicotine and cisplatin. There are always happen in adenocarcinoma or squamous cancer cells. Intriguingly, we determine nicotine induces NSCLC cancer cell lines proliferation, cell growth, and migration using immunoblot and immunofluorescence analysis. Nicotine may binds to nicotinic-acetylcholine receptors (α7nAchR) or through EGFR leading to activation of downstream effects, such as cancer cell growth. Results showed us that nicotine induced through α7nAchR, but not EGFR, in the cytosol not in the nucleus. Both of α7nAchR and EGFR in A549 cells are increased after HDAC1^+/+^ transfection in the cytosol and in the nucleus (Fig. 5B). In particular, E2F was increased by nicotine in the cytosol and in the nucleus. After nicotine, A549 cells nucleus were observed cisplatin-resistance by E2F. Groundbreaking, we observed pRb decreases in the nucleus, but after with nicotine, pRb increases in the nucleus and observed cisplatin-resistance effect (Fig. 5C). Also of concern, α7nAchR and EGFR only observed in cell membrane after nicotine addition, cytosol and nucleus did not found them (Fig. 5D). In the nucleus location, we found that HDAC1 and E2F binding together, and HDAC1 and Rb using immunoprecipitation analysis (Fig. 5E). When cisplatin binds with DNA in A549 cells without nicotine, we could find HDAC1, E2F and Rb binding together in the nucleus. After nicotine with squamous carcinoma cell (H157), we found HDAC1, Rb, and E2F were observed in the cytosol. In the nucleus, HDAC1 was increased by HDAC1^+/+^ transfection compared with control. In contrast, Rb and E2F were no effects changes (Fig. 5F). The protein of Rb and E2F was observed by Cisplatin in the cytosol and nucleus, but pRb was only observed in the cytosol, not in the nucleus. After with nicotine in H157 cell, pRb was observed in nucleus control and HDAC1^+/+^, but did not find in the cytosol. The protein of E2F with nicotine was increased by HDAC1^+/+^ transfection, but no change in the nucleus compared with nucleus control. Cisplatin-resistance effects was observed with nicotine treated H157 cell. E2F by cisplatin was not change in the cytosol, but decreased was in the nucleus (Fig. 5G). That we think cisplatin was released by nicotine.

### Nicotine induced cisplatin-resistance activity through regulation of Cyclin E/E2F and p53/p21/p27 in H157 cell

On the other hand, at high nicotine concentration (> 1 μM) with consistent cytotoxic effects and appeared to be due to direct cell kill. The concentrations of nicotine promote cell proliferation correspond to the low concentrations, while high concentrations are cytotoxic. As is well known, lung cancer remains one of the most common types of fatal malignancies. Nicotine could induce the proliferation of a variety of lung carcinoma cell clines, but there is no evidence that nicotine itself provokes cancer. We found H157 cell viability was increased at 1.0 μM and 2.0 μM nicotine concentrations in H157 squamous cancer cell. Time different at 1 μM was determine cell viability (%) using MTT assay. At 72 h, we found to have higher cell viability (%) (Fig. 6A). Furthermore, we detected HDAC1 inhibitor, VPA, and cisplatin-related with short-(24 h) and long-term (48 h) nicotine treatment in H157 cells. Cisplatin resistance no found at 24 and 48 h nicotine treatment. VPA, cisplatin and their combination have significantly different cell viability (%) at 24 and 48 h nicotine treatment, when compared with their own control (Fig. 6B). Therefore, we determine nicotine treatment at 72 h induced cisplatin resistance related with cell cycle regulation in H157 cell by western blot. First, we want to know cisplatin and nicotine related with HDAC1 in the cytosol or in the nucleus using immunofluorescent staining. Result showed HDAC1 fluorescent positive cells increases after cisplatin in the nucleus, but after nicotine in the nucleus, HDAC1 fluorescent positive cells increases in the cytosol (Fig. 6C). Nicotine induced cancer cell migration at 24 and 48 h (Fig. 6D). Nicotine induced cisplatin resistance through α7 nicotinic-acetylcholine receptors (α7nAChR) and EGFR. From Fig. 6E results showed without nicotine, protein of HDAC1, α7 nicotinic-acetylcholine receptors (α7nAChR) and EGFR has VPA, cisplatin and their combination by western blot analysis was decreased compared with control, however, after with nicotine, VPA, cisplatin and their combination no effects on these protein decreases. In H157 cells without nicotine, cell cycle check proteins, Rb/2F/Cyclin E/Cyclin D1, have decreased by cisplatin and VPA combined with cisplatin, once nicotine add, protein of Rb/2F/Cyclin E/Cyclin D1 was lost function by cisplatin (Fig. 6F).

So far encouraging results from cell cycle regulation, cisplatin and VPA still have good functions in spite of with or without nicotine. Given these prerequistes and a receptivity for cell cycle regulation, suppression genes, p21, p27 and p53 were increased by VPA, cisplatin and their combination, after nicotine treatment, p21 and p53 have a little lower than without nicotine, but VPA combined with cisplatin has a significantly decreased. Cisplatin resistance was observed in p27 suppression gene. We found p27 protein no significant change.

### Nicotine induced cisplatin-resistance activity in H520 cell

To check receptors, α7nAChR and EGFR, in the cytosol expression by western blotting analysis. Results showed α7nAChR and EGFR expression in nicotine treatment, HDAC1 plasmid transfect (HDAC1^+/+^) and HDAC1 inhibitor, VPA, in the cytosol were observed increases and decreases regulation (Fig. 7A). Contractually, nicotine induced squamous cancer cells H520 growth, in spite of HDAC1 or not. Cell cycle regulation could be found it’s expression levels in HDAC1^+/+^ plasmid transfection, HDAC1 inhibitor (VPA). Cell cycle related proteins, Cyclin A/ Cyclin D/ Cyclin E/ Cyclin B/E2F increases and p53/p27/p21/pRb decreases, were found by HDAC1. Nicotine treatment led HDAC1^+/+^ and VPA loss function (Fig. 7B). When the extent of damage is limited, cisplatin adducts induce an arrest in the S and G2 phases of the cell cycle. Cisplatin resistance was observed regulation by nicotine (Fig. 7C). Cisplatin activates in the cytoplasm intracellularly spontaneously. Cyclin-dependent kinase inhibitors, p53/p21/p27, results in cell cycle arrest and apoptosis occurs cisplatin resistance (Fig. 7D).

## Discussion

Chemotherapy drug, cisplatin, target NSCLC cells at different phases of the process of forming the cell cycle to interfere cells in various phases of the cell cycle. A low concentration of Chloride ions resulted in the generation of mono- and bi-cisplatin forms [17,18]. In the cytoplasmic cisplatin has the potential to deplete reduced equivalents and to destroy the redox balance toward oxidative stress [19,20]. In the nucleus, Platinum-based doublet chemotherapy remains the mainstay for advanced NSCLC. Cisplatin, a highly effective and widely used chemotherapeutic agent [21]. Cisplatin can block the synthesis of DNA, thereby blocking the division of cancer cells, is a non-specific choice of broad-spectrum anti-cancer drugs, commonly used in the treatment of NSCLC lung cancer [22]. Several mechanisms account for the cisplatin-resistant of tumor cells have been described. Some reports elucidated cancer cells growth may induce by HDAC1 factor to regulation [23,24]. Histone deacetylases (HDAC) modulates acetylation of lysine residues, formation of the core histone complex (H2A, H2B, H3 and H4) plays a crical role to regulate cell cycle progression [25,26]. HDAC2 was dispensable for HDAC1 binding to HDAC2-activated targets, HDAC2 was required for the recruitment of HDAC1 to repressed HDAC2 gene (Fig. 1S). Cisplatin is often selected due to in case of stage II and stage III disease following surgery for localized non-small-cell lung cancer (NSCLC) [27]. Cisplatin and HDAC1 bound suppressed E2F and Rb transcription into the cytosol resulted in cell cycle stop. These results could find from Fig. 2, Fig. 3, Fig. 4, and Fig. 5 supported. Nicotine and HDAC1 protein interaction expression in A549 adenocarcinoma cytosol and nuclear (Fig. 5). The expression of HDAC1 was found to be associated with most cell types of NSCLC (Fig. 6 and Fig. 7). Our preliminary results showed A549 adenocarcinoma may response nicotine effects resulted in cisplatin resistance. At low concentration, nicotine induced squamous cancer cell growth, in spite of chemotherapy drugs or not (Fig. 6A and Fig. 6B). Some reports elucidated cancer cells growth may induce by nicotine and HDAC1 regulation. Nicotine binds to nicotinic-acetylcholine receptors (α7nAchR) and EGFR receptor leading to activation of the HDAC1 signaling pathway (Fig. 6E and Fig. 7A). Nicotine increases migration and invasion of lung cancer cells through activation of the α7nAChR (Fig. 6C and Fig. 6D). The concentration of nicotine stimulated cell growth correspond to low concentration was needed, while high concentration was cytotoxic [28]. Nicotine also stimulates angiogenesis and tumor growth which is also mediated through nicotinic-acetylcholine receptors (nAChR), possibly involving endothelial growth factor (EGF). Nicotine is significantly higher cisplatin resistance in NSCLC (Fig. S3). During nicotine in concentrations as low as 1 μM, nicotine activates cell migration, proliferation, survival, and anti-apoptotic effects exerted modulation chemotherapeutics on several different malignant cell lines (Fig. 6 and Fig. S4). Cisplatin anticancer effects by multiple mechanisms, despite a consistent rate of initial responses into nucleus, nicotine via these mechanisms lead to the development of chemoresistance resulted in therapeutic failure (Fig. 8). Cisplatin involves the DNA-damage response is time- and dose-dependant [29,30]. Cisplatin involves the DNA-damage response is time- and dose-dependant [31]. Nicotine can prevent apoptosis induced by various agents in NSCLC. Cell growth inhibition and apoptosis is a result of HDAC1-mediated cyclin D/E suppression. Down-regulation of HDAC1 expression mediates proliferation inhibition and cell cycle arrest [32,33]. Therefore, HDACs causes the inhibition of transcription activation of cancer growth. As in some clinical settings cisplatin constitutes the major therapeutic option, the development of chemosensitization strategies constitute a goal with important clinical implications

## Methods

### Cell culture

NSCLC lung cancer lines, adenocarcinoma (A549) and squamous cell carcinoma (H157 and H520) were obtained from human lung epithelial cells. A549 (BCRC number 60074) and NCI-H520 (BCRC number 60124) lung cancer cells were purchased from the Bioresource Collection and Research Center (BCRC) in Taiwan. H157 (ECACC no. 07030901) was purchased from <u>CellBank Australia</u>. Adenocarcinoma lung cancer cells, A549, was grown in 90% Ham’s F12K medium with 2 mM L-glutamine adjusted to contain 1.5 g/L sodium bicarbonate, 10 mM HEPES, 1 mM sodium pyruvate, 4500 mg/L glucose, and 10% fetal bovine serum for use in incubators using 5% CO_2_ in air. Squamous cell carcinoma, NCI-H520, 90% RPMI 1640 medium and 10% fetal bovine serum. NCI-H157 was grown in the DMEM:HAMS F12 (1:1), 2 mM Glutamine, 10% Foetal Bovine Serum (FBS) and 0.5 ug/ml sodium hydrocortisone succinate. Lung cancer cells were grown in F-12, EMEM, Dulbecco’s Modified Eagle Medium (DMEM) and RPMI-1640 medium, respectively, with 100 units/mL penicillin, and 100 μg/mL streptomycin, 5% CO_2_ atmosphere at 37 °C.

### Cell culture with or without nicotine treatment and transfection HDAC1^+/+^, siRNA plasmid, Valproic acid (VPA) and Cisplatin (Cis)

Adenocarcinoma cells (A549) and squamous cancer cells (H157 and H520) lung cancer cell lines were maintained in continuous culture at 37°C, 5% CO2 in medium containing penicillin and streptomycin mixture, L-glutamine, sodium bicarbonate, and 10% fetal bovine serum (FBS) without or with nicotine (1μM) grown in 6-well plate. Cells were treated with HDAC1 plasmid (HDAC1^+/+^, 3 ug/ml) transfected to cells using Lipofectamine(tm) 2000 according to the manufacturer’s instructions. Cells were inhibited HDAC1 expression treated with 1 mM valproic acid (VPA; a specific inhibitor of HDAC1). Cisplatin, a chemotherapy drug (Cis, 25 μM), treated lung cancer cell lines 24 hours with or without nicotine to establish cisplatin-resistance cell lines. To establish HDAC1 knockdown cell lines, HDAC1 control siRNA lentivirus plasmid-A, were respectively after transfection for 24 h. HDAC1siRNA plasmid (1ug/ml) will be transiently transfected into lung cancer cells. HDAC1 small interfering RNAs (HDAC1 siRNAs): 5^”^-CCCAUAACUUGCUGUUAAA-3^’^ (Santa Cruz Biotechnology, Inc.) using Lipofectamine 2000 reagent (Invitrogen), then harvested for further analysis 24 h after transfection. For nicotine treatment, all lung cells received 1 uM nicotine for 24 h.

### MTT assay (Cell viability)

Lung adenocarcinoma (A549) growth rate were determined by MTT assay. A549 cells (1×10^6^ cells/well) were seeded into 96-well culture plates. A549 cells were treated with HDAC1 plasmid (HDAC1^+/+^), Valproic acid (VPA, 1 mM), and Cisplatin (Cis, 25 μM) without nicotine treatment in Figure 1A result. Cisplatin (Cis) was treated with 5, 10, 25 and 50 μM in A549 cell to determine cell viability without nicotine in Figure 1B result. Valproic acid (VPA) was treated with 0.6, 12.5, 25, 50 and 100 μM in A549 cell to determine cell viability without nicotine in Figure 1C result. For nicotine treatment, H157 lung cells received 0.1, 0.5, 1.0, 1.5, 2.0, 2.5, 3.0 μM nicotine for 24 h in Figure 6A result, and incubated with 1 μM nicotine (Sigma, Missouri, USA) for H157 for 6, 12, 24, 72, 96 and 120 h in the Figure 6B result. H157 lung cells incubated with 1 μM nicotine for 24 h as short-term, for 72 h as as long-term in the Figure 6C result. An MTT solution (5 mg/ml, 1ml) was added to 9 ml RPMI 1640 media supplemented with 10% FBS. After added PMS (5 mg/ml, 10 ul) into solution and mix. Mix solution (100 ul) into each well and the plates were incubated at 37°C for 2 h. After centrifugation at 3,000 rpm for 10 min, remove the supernatant. The remaining formazan pellet was dissolved completely in DMSO (100 ul). An ELISA plate reader was used to measure the absorbance at 570 nm to determine the amount of pellet.

### Cell migration Assay

A549 and H157 cells to adhere and spread completely. Plate H157 cells to create a confluent monolayer onto the prepared 60 mm dish; create a wound by manually scraping the cell monolayer with a p200 pipet tip. Wash the cells once with 1 mL of desired medium and replace with 1mL of the RPMI 1640 medium. The wound should be created relative to the marking/reference point on the dish. Acquire the first image by using the markings on the culture dish as a reference point. Incubate dishes in a culture incubator for 24-48 h.

### Immunofluorescence staining

Lung cancer cells, A549 and H157, incubated with HDAC1 and NF-κB primary antibody at 4°C overnight and Texas Red-conjugated secondary antibody and for 2 h at room temperature. Extensive washing with PBS will be performed between each step and before mounting with Fluoromount G and then examined by fluorescence microscope (Zeiss Axioskop2 fluorecent microscope). Match the photographed region acquired in first image and second image.

### Western blot analysis

A549 lung cancer cell protein are collected and dissolved in lysis buffer (10 mM Tris-HCl pH 7.4, 100 mM NaCl, 1 mM EDTA, 1Mm EGTA, 1mM NaF, 2mM Na3VO4, 1% Triton X-100, 10% glycerol, 0.1% SDS, 0.5% deoxycholate, 1 mM PMSF, 1mg/ml leupeptin, and 5ml/ml aprotinin). The extracts will be centrifuged at 13,000 rpm for 15 min at 4°C. The protein concentration in supernatant will be determined by Bradford assay using the Bio-Rad Protein Assay kit (Bio-Rad, Hercules, CA, USA). Equal amounts of total protein are separated on 8∼12% SDS–PAGE gels, then transferred on to PVDF membrane with transfer system cell (Bio-Rad), incubated with 5% no-fat milk and then hybridized with α7nAchR, EGFR, Cyclin D/E/B/A, HDAC1/2/3/4, E2F/pRb/Rb, p53/p27/p21, Caspase 3/Bax, NF-KB/γ-H2AX/ATM, GAPDH/α-tubulin/β-actin primary antibody (Santa Cruz Lab, CA, USA), and then recognized with the secondary antibody. Bands development were presented in ECL chemiluminescence detection system (Amersham Intl., Buckinghamshire, UK).

### Immunoprecipitation analysis

The A549 cells will be collected and dissolved in EBC buffer (50 mM Tris pH7.6, 120 mM NaCl, 0.5% Nonidet P-40, 1 mM EDTA, 1 mM β-mercaptoethanol, 50 mM NaF, 1 mM Na3VO4, plus protease inhibitors). After determination of the concentration, 40μg of protein lysates will be incubated with anti-HDAC1 protein (from Santa Cruz, USA) followed by protein G-PLUS agarose precipitation. The immunoprecipitates will be separated by SDS-PAGE and the above-mentioned E2F and Rb proteins, after transfer from gel to nitrocellulose membrane, will be detected by anti-secondary antibody (from Santa Cruz).

### Statistical analysis

Quantitative all the results were shown as the mean ± SEM. The statistical significance of difference between means was evaluated using one-way analysis of variance (ANOVA), followed by post hoc analysis as comparison tests. *P*<0.05, *P*<0.01 was regarded as statistically significant.

## Acknowledgements

This work was supported by grants from Ministry of Science and Technology (MOST 104-2811-B-009-007, MOST 105-2811-B-039-008 and MOST 106-2811-B-650-003).

## Authors’ contributions

CY Peng designed the experiments and analyzed the data. JP Wu performed the experiments and wrote the paper.

## Figure Legends

**Fig. 5. Regulation of A549 and H157 cells growth E2F/Rb signaling with or without nicotine in the cytosol and nucleus**. (A). HDAC1 regulated A549 cells growth with or without nicotine in the cytosol and nucleus. HDAC1^+/+^, HDAC1 plasmid transfection; Cis, cisplatin. (B). Nicotine induced α7nAChR, EGFR and E2F related with HDAC1 in A549 cells cytosol and nucleus. (C) In the nucleus, pRb, Rb and E2F are exchanged each other with cisplatin resistance in A549 cells with or without nicotine using western blotting. HDAC1^+/+^, HDAC1 plasmid transfection; Cis, cisplatin. (D). Location of α7nAChR and EGFR in the cell membrane in A549 cells with or without nicotine. (E). HDAC1/E2F and HDAC1/Rb bind together using immunoprecipitation analysis. (F). HDAC1/Rb/E2F regulated H157 cells growth with or without nicotine in the cytosol and nucleus. HDAC1^+/+^, HDAC1 plasmid transfection. (G). HDAC1/pRb/E2F regulated H157 cells cisplatin resistance with or without nicotine in the cytosol and nucleus. HDAC1^+/+^, HDAC1 plasmid transfection; Cis, cisplatin.

**Fig. 6. Establish nicotine-induced H157 cells cisplatin resistance through HDAC1 regulated E2F/Rb model**. (A) Dose- and time-dependent of H157 nicotine treatment cell viability (%) by MTT assay. (B). Short- and long-term nicotine treatment for VPA and Cis. Cis, cisplatin; VPA, valproic acid. (C). Immunofluorescence assay of HDAC1 protein and H33256 in H157 cells treated with nicotine (1μM) and Cis for location analysis. Cis, cisplatin. (D). Cell migration of H157 squamous lung cancer cells with nicotine at 24 and 48 h. (E). Western blotting of membrane receptors, EGFR, α7nAChR and HDAC1 related with VPA and Cis treatment to determine nicotine lead cisplatin resistance. Cis, cisplatin; VPA, valproic acid. (F). Western blotting of cell cycle related proteins including Rb/E2F/Cyclin E/Cyclin D/Cyclin A/Cyclin B and p53/p21/p27 associated with HDAC1 and Cis in H157 with or without nicotine. Cis, cisplatin; VPA, valproic acid.

**Fig. 7. Nicotine induced H520 cells cisplatin resistance through HDAC1 regulated E2F/Rb**. (A).Western blotting of membrane receptors, EGFR, α7nAChR and HDAC1 related with VPA and HDAC1^+/+^ treatment to determine nicotine lead cell growth. HDAC1^+/+^, HDAC1 plasmid transfection; VPA, valproic acid. (B). Western blotting of cell cycle related proteins, Cyclin A/Cyclin D/Cyclin E/E2F/Cyclin B and p53/p21/p27/pRb in H520 with or without nicotine treatment lead cell growth. HDAC1^+/+^, HDAC1 plasmid transfection; VPA, valproic acid. (C). Western blotting of cell cycle related proteins, HDAC1/HDAC2/pRb/Cyclin E/E2F/Cyclin A/Cyclin B/Cyclin D in H520 cells with or without nicotine treatment lead cell growth and cisplatin resistance. HDAC1^+/+^, HDAC1 plasmid transfection; VPA, valproic acid; Cis, cisplatin. (E). Western blotting of cell cycle suppressor proteins, p53/p21/p27 in H520 cells with or without nicotine treatment lead cell growth and cisplatin resistance. HDAC1^+/+^, HDAC1 plasmid transfection; VPA, valproic acid; Cis, cisplatin.

**Fig. 8. Schematic diagram of cross talk between HDAC1 and cell cycle signaling pathways**. The molecular mechanisms account for the nicotine induces cisplatin-resistant phenotype. In the nucleus, nucleophilic cisplatin insert and binds DNA tightly due to with activation of the generation of DNA damage responses and the induction of apoptosis. HDAC1 affecting off-target cell cycle E2F/Rb induces growth by cisplatin, however, nicotine induces cisplatin resistance molecular circuitries is because of present obvious links with loss cisplatin-elicited signals. Nicotine via these mechanisms lead to the development of chemoresistance resulted in therapeutic failure.

